# Estimates of cortical column orientation improve MEG source inversion

**DOI:** 10.1101/810267

**Authors:** James J Bonaiuto, Fardin Afdideh, Maxime Ferez, Konrad Wagstyl, Jérémie Mattout, Mathilde Bonnefond, Gareth R Barnes, Sven Bestmann

**Affiliations:** Institut des Sciences Cognitives Marc Jeannerod, CNRS UMR 5229, Bron, France; Université Claude Bernard Lyon 1, Université de Lyon, France; Lyon Neuroscience Research Center, Brain Dynamics and Cognition team, INSERM UMRS 1028, CNRS UMR 5292, Bron, France; University of Cambridge, Department of Psychiatry, Cambridge CB2 0SZ, UK; Wellcome Centre for Human Neuroimaging, UCL Queen Square Institute of Neurology, University College London (UCL), London, WC1N 3BG, UK; Dept of Clinical and Movement Neuroscience, UCL Queen Square Institute of Neurology, University College London (UCL), London, WC1N 3BG, UK

**Keywords:** source inversion, dipole orientation, cortical columns, cortical surface, high precision MEG

## Abstract

Determining the anatomical source of brain activity non-invasively measured from EEG or MEG sensors is challenging. In order to simplify the source localization problem, many techniques introduce the assumption that current sources lie on the cortical surface. Another common assumption is that this current flow is orthogonal to the cortical surface, thereby approximating the orientation of cortical columns. However, it is not clear which cortical surface to use to define the current source locations, and normal vectors computed from a single cortical surface may not be the best approximation to the orientation of cortical columns. We compared three different surface location priors and five different approaches for estimating dipole vector orientation, both in simulations and visual and motor evoked MEG responses. We show that models with source locations on the white matter surface and using methods based on establishing correspondences between white matter and pial cortical surfaces dramatically outperform models with source locations on the pial or combined pial/white surfaces and which use methods based on the geometry of a single cortical surface in fitting evoked visual and motor responses. These methods can be easily implemented and adopted in most M/EEG analysis pipelines, with the potential to significantly improve source localization of evoked responses.

## Introduction

Non-invasive measures of brain activity such as magnetoencephalography (MEG) and electroencephalography (EEG) are powerful tools for generating insights into human brain function with millisecond-scale temporal resolution. However, determining the current distribution that gives rise to the signals measured from EEG and MEG sensors is challenging (Baillet et al., 2001; Darvas et al., 2004; Fukushima et al., 2012; Haufe et al., 2011; Mattout et al., 2006). In order to simplify the source localization problem, many techniques introduce constraints to the dimensionality of source space. These constraints embody assumptions about how the brain generates the signals which we can measure from outside of the head.

One of these assumptions is that signals measured by M/EEG sensors are predominantly generated by large pyramidal neurons in deep cortical layers, which are arranged in parallel columns so that their cumulative activity produces a measurable extracranial signal (Baillet, 2017; Buzsáki et al., 2012; Murakami and Okada, 2006; Okada et al., 1997). Two commonly used source localization constraints based on this assumption are that the locations of source dipoles are restricted to locations on a mesh of the white matter surface as is it is closest to the deep cortical layers (Dale and Sereno, 1993; Henson et al., 2009; Hillebrand and Barnes, 2003, 2002; Mattout et al., 2007), and that the orientation of dipoles is orthogonal to this surface (Hämäläinen and Ilmoniemi, 1984, 1994; Henson et al., 2009; Hillebrand and Barnes, 2003; Lin et al., 2006; Salmelin et al., 1995), thereby approximating the orientation of cortical columns (Nunez and Srinivasan, 2006; Okada et al., 1997).

Using vectors orthogonal to the cortical surface may not be the best approximation to the orientation of cortical columns. Cortical folding patterns may result in curved cortical columns, and therefore their orientation with respect to the cortical surface could be different along the gray / white matter (white matter surface) and CSF / gray matter (pial surface) boundaries. Moreover, induced activity in low and high frequency bands can predominate in deep or superficial cortical layers (Bastos et al., 2015; Bonaiuto et al., 2018a; Buffalo et al., 2011; Haegens et al., 2015; Maier et al., 2010; Spaak et al., 2012; van Kerkoerle et al., 2014), and therefore the white matter surface may not be the optimal source location model. In the past, however, the contribution of inaccuracies in dipole location and orientation constraints to source localization error has likely been insignificant in the face of within-session participant movement, co-registration error, and the relatively low resolution of cortical surface reconstructions. However, the recent development of techniques for high precision MEG (Bonaiuto et al., 2018b, 2018a; Meyer et al., 2017; Troebinger et al., 2014b, 2014a) allow us to compare competing current-flow orientation models in more detail.

Here, we set out to determine a better way to estimate the location and orientation of source dipoles based on MRI-derived cortical surfaces. We tested three different cortical surfaces for determining dipole locations: 1) white matter, 2) pial, and 3) combined white matter/pial, and five different methods for computing dipole orientations: 1) downsampled surface normals, 2) cortical patch statistics, 3) original surface normals, 4) link vectors, and 5) variational vector fields. The most commonly used method, downsampled surface normals (Dale and Sereno, 1993; Fuchs et al., 1994; Hämäläinen and Hari, 2002; Hillebrand and Barnes, 2003; Lin et al., 2006), involves downsampling (decimating) the original cortical surface, and then computing the normal vector at each vertex as the mean of the normal vectors of each surface face it is connected to. While surface decimation increases the computational tractability of source inversion, it distorts the surface faces and therefore biases the surface normal vector estimates. The cortical patch statistics method was therefore designed to compute normal vectors by averaging the individual normal vectors from vertices adjacent to the nearest vertex in the original (down-sampled) mesh (Lin et al., 2006). The original surface normals method takes advantage of the fact that the surface decimation algorithm used here maintains a correspondence between the downsampled and original surface meshes, and uses the normal vectors of the corresponding vertices from the original cortical surface. These three methods involve computation of dipole orientation based on the geometry of a single cortical mesh: the white matter *or* pial surface. In contrast, the link vectors (Dale et al., 1999) and variational vector field (Fischl and Sereno, 2018) approaches establish correspondences between the white matter and pial surface meshes. The link vectors approach simply uses the vectors connecting each vertex on the white matter surface with the corresponding vertex on the pial surface (Dale et al., 1999). The variational vector field method constructs a field of correspondence vectors between the original white matter and pial surfaces which are constrained to be approximately normal to each cortical surface and parallel to each other (Fischl and Sereno, 2018).

We first compared the resulting orientation vectors from each method in terms of the angular difference at each surface vertex. We then ran simulations of single dipoles at a given orientation, and subsequently performed source reconstruction using various dipole orientations, noise levels, and co-registration error magnitudes. Finally, we compared the methods using evoked visual and motor responses in MEG data from human participants.

## Methods

Data from eight healthy, right-handed, volunteers with normal or corrected-to-normal vision and no history of neurological or psychiatric disorders was used for our analyses (six male, aged 28.5 ± 8.52 years; Bonaiuto et al., 2018a; Little et al., 2018). The study protocol was in accordance with the Declaration of Helsinki, and all participants gave written informed consent which was approved by the UCL Research Ethics Committee (reference number 5833/001). All analysis code is available at https://github.com/jbonaiuto/dipole_orientation.

### MRI acquisition

Prior to MEG scanning, two MRI scans were acquired with a 3T whole body MR system (Magnetom TIM Trio, Siemens Healthcare, Erlangen, Germany) using the body coil for radio-frequency (RF) transmission and a standard 32-channel RF head coil for reception. The first was a standard T1 for individual head-cast creation (Meyer et al., 2017), and the other was a high resolution, quantitative multiple parameter map (MPM; Weiskopf et al., 2013) for MEG source location.

The first protocol used a T1-weighted 3D spoiled fast low angle shot (FLASH) sequence with 1 mm isotropic image resolution, field-of view set to 256, 256, and 192 mm along the phase (anterior-posterior, A–P), read (head-foot, H–F), and partition (right-left, R–L) directions, respectively. The repetition time was 7.96 ms and the excitation flip angle was 12°. After each excitation, a single echo was acquired to yield a single anatomical image. A high readout bandwidth (425 Hz/pixel) was used to preserve brain morphology and no significant geometric distortions were observed in the images. Acquisition time was 3min 42s. A 12 channel head coil was used for signal reception without using either padding or headphones.

The second, MPM, protocol consisted of acquisition of three differentially-weighted, RF and gradient spoiled, multi-echo 3D fast low angle shot (FLASH) acquisitions and two additional calibration sequences to correct for inhomogeneities in the RF transmit field (Callaghan et al., 2015; Lutti et al., 2012, 2010), with whole-brain coverage at 800 μm isotropic resolution.

The FLASH acquisitions had predominantly proton density (PD), T1 or magnetization transfer saturation (MT) weighting. The flip angle was 6° for the PD- and MT-weighted volumes and 21° for the T1 weighted acquisition. MT-weighting was achieved through the application of a Gaussian RF pulse 2 kHz off resonance with 4 ms duration and a nominal flip angle of 220° prior to each excitation. The field of view was 256 mm head-foot, 224 mm anterior-posterior (AP), and 179 mm right-left (RL). Gradient echoes were acquired with alternating readout gradient polarity at eight equidistant echo times ranging from 2.34 to 18.44 ms in steps of 2.30 ms using a readout bandwidth of 488 Hz/pixel. Only six echoes were acquired for the MT-weighted acquisition in order to maintain a repetition time (TR) of 25 ms for all FLASH volumes. To accelerate the data acquisition, partially parallel imaging using the GRAPPA algorithm was employed with a speed-up factor of 2 in each phase-encoded direction (AP and RL) with forty integrated reference lines.

To maximize the accuracy of the measurements, inhomogeneity in the transmit field was mapped by acquiring spin echoes and stimulated echoes across a range of nominal flip angles following the approach described in Lutti et al. (2010), including correcting for geometric distortions of the EPI data due to B0 field inhomogeneity. Total acquisition time for all MRI scans was less than 30 min.

Quantitative maps of proton density (PD), longitudinal relaxation rate (R1 = 1/T1), MT and effective transverse relaxation rate (R2* = 1/T2*) were subsequently calculated according to the procedure described in Weiskopf et al. (2013).

### FreeSurfer surface extraction

FreeSurfer (v5.3.0; Fischl, 2012) was used to extract cortical surfaces from the MPMs for MEG source localization. We used a custom FreeSurfer surface reconstruction procedure to process MPM volumes, using the PD and T1 volumes as inputs (Carey et al., 2017), resulting in surface meshes representing the pial surface (adjacent to the cerebro-spinal fluid, CSF), and the white/grey matter boundary (**Figure 1**). FreeSurfer creates the pial surface by expanding the white matter surface outward to the cortex/CSF boundary. This is done by minimizing an energy functional which includes terms promoting surface smoothness and regularity as well as an intensity-based term designed to determine the cortex/CSF boundary based on local volume intensity contrast (Dale et al., 1999). Because this process involves moving the vertices of the white matter surface based on the gradient of the energy functional, the result is a one-to-one correspondence between white matter and pial surface vertices. We used a custom routine to downsample each of these surfaces by a factor of 10 while maintaining this correspondence. This involved using MATLAB’s reducepatch function to remove vertices from, and re-tesselate the pial surface, and then removing the same vertices from the white matter surface and copying the edge structure from the pial surface. This yielded two meshes of the same size (same number of vertices and edges), comprising about 30,000 vertices each (M = 30,094.75, SD = 2,665.45 over participants).

### Dipole orientation computation methods

The downsampled surface normal method computes, at each vertex of the decimated mesh, the average of the normal vectors of each adjacent face (**Figure 1**). This method was implemented using the spm_mesh_normals function in SPM12. The cortical patch statistics method computes, at each vertex of the decimated mesh, the average surface normal vector over all vertices in the original, non-decimated mesh which are adjacent to the corresponding original mesh vertex (**Figure 1**). The original surface normal method is implemented in the same way as the downsampled surface normals method, but is applied to the original, non-decimated mesh (**Figure 1**). Because our decimation procedure only removed vertices from the original surface, the resulting vectors can then be mapped onto the vertices of the decimated mesh used for source localization. The link vectors method takes advantage of the fact that our decimation routine maintains the correspondence between white matter and pial surface vertices, and for each vertex on the pial surface, uses the vector linking it to the corresponding vertex on the white matter surface (**v_i_**=**w_i_** - **p_i_**, for the *i*^th^ white matter vertex, *w_i_* and pial vertex p_i_; **Figure 1**). The variational vector field method constructs a vector field linking the white matter and pial surfaces, by using gradient descent to minimize an energy functional that encourages vectors to approximate surface normals and to be parallel to each other (Fischl and Sereno, 2018) (**Figure 1**). The angular difference between any two vectors, v1 and v2, was computed using the formula: atan2(|| **v_1_** × **v_2_** ||, **v_1_** · **v_2_**).

Vectors obtained from each method were used to construct the lead field matrix of the forward model used for source inversion of the simulated or experimental data. The models constructed using each method were compared to each other based on relative Bayesian model evidence, as approximated by differences in free energy:

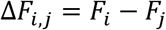

where *F_i_* and *F_j_* are the free energy values of models *i* and *j*, respectively. Free energy is a parametric metric rewards fit accuracy and penalizes model complexity (Bonaiuto et al., 2018b; Friston et al., 2008, 2007; Henson et al., 2009; López et al., 2014; Wipf and Nagarajan, 2009):

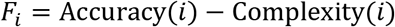

The first term is the log model evidence: the log of the probability of the data, given the model and parameters, and the second term is the Kullback-Liebler divergence between the true posterior density and an approximate posterior density. Because the second term is always positive, free energy provides a lower bound on the model evidence (Penny et al., 2010).

The best overall dipole orientation method and source space surface model was determined using random effects family level Bayesian inference (Penny et al., 2010) as implemented by the spm_compare_families method in SPM12. This method groups models based on visual ERF 1 and 2 and the motor ERF in all participants into ‘families’, and then combines the evidence of models from the same family and computes the exceedance probability for each family. The exceedance probability corresponds to the belief that a particular model family is more likely than the other model families tested, given the data from all participants. We first grouped models into families based on dipole orientation method / source space surface model combinations (e.g. downsampled surface normals / pial surface) to determine the best combination over all ERFs and participants. We then grouped models based on dipole orientation method, and finally based on source space surface model.

**Figure 1.**
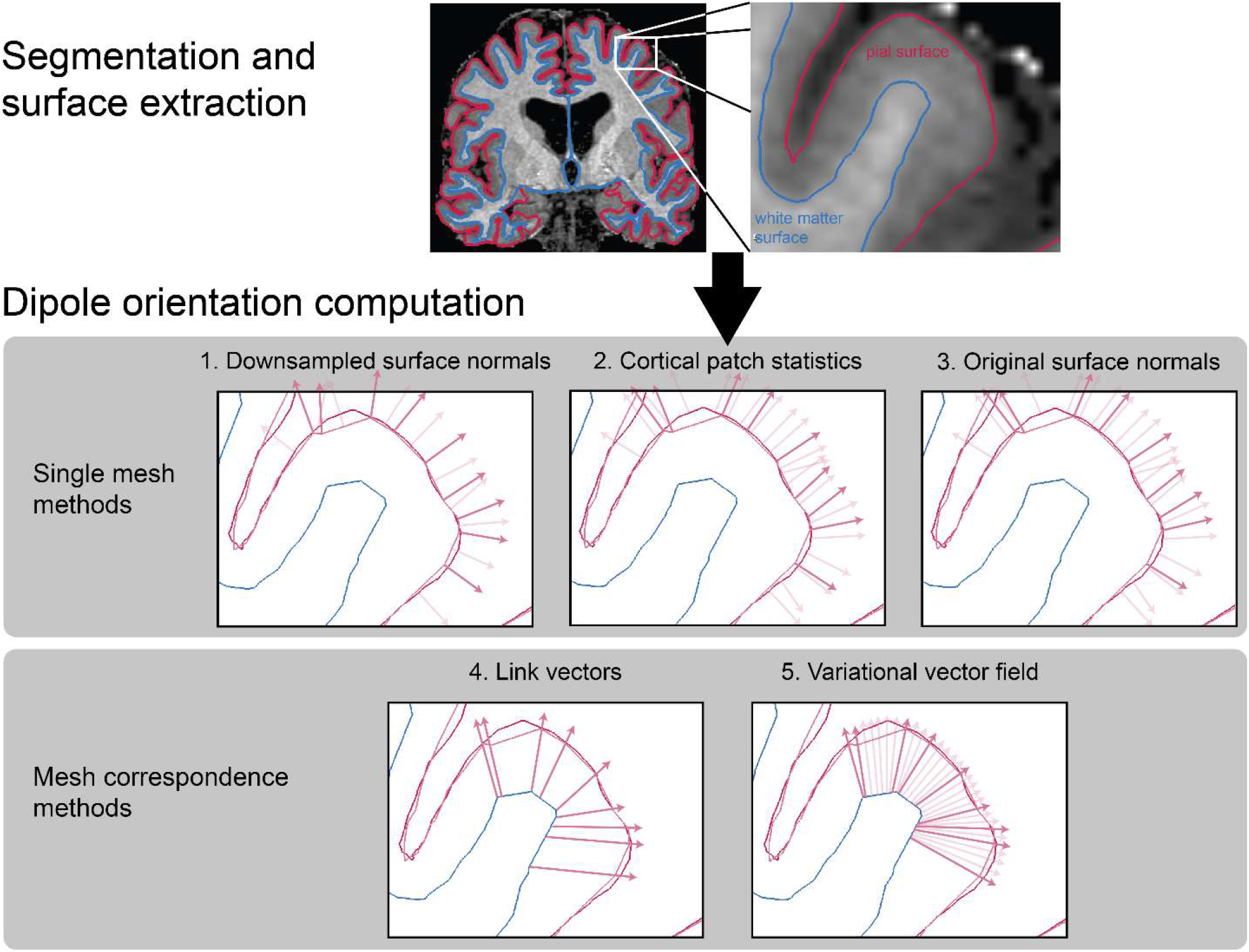
Dipole orientation models. Pial and white matter surfaces are extracted from proton density and T1 weighted quantitative maps obtained from a multi-parameter mapping MRI protocol. Dipole orientation vectors are computed from these surfaces using five different methods. The downsampled surface normal and original surface normal methods compute vectors at each vertex (dark red) as the mean of the normal vectors of the surface faces they are connected to (light red). The cortical patch statistics method computes the mean of the normal vertices adjacent to the corresponding vertices in the original mesh. The link vectors method computes vectors which link corresponding vertices on the white matter and pial surfaces. The variational vector field method constructs a field of vectors which are approximately parallel to each other and orthogonal to the pial surface (shown in light red for the original surfaces and dark red for the subset of vertices in the downsampled surfaces).

### Simulations

All simulations were based on a single dataset acquired from one human participant. This dataset was only used to determine the sensor layout, sampling rate (1200 Hz, downsampled to 250 Hz), number of trials (515), and number of samples (251) for the simulations. All simulations and analyses were implemented using the SPM12 software package (http://www.fil.ion.ucl.ac.uk/spm/software/spm12/).

In each simulation, we specified spatially distributed source activity centered at a single vertex on the pial surface. We simulated a Gaussian activity time course in this vertex, centered within the epoch, with a width of 25ms and a magnitude of 10nAm. We then spatially smoothed this simulated dipole time course with a Gaussian kernel (FWHM=5mm), to obtain a patch of spatially distributed activity. Within this patch, the orientation of each vertex differed, but was specified by the same rule using the link vectors method. We then used a single shell forward model (Nolte, 2003) to generate a synthetic dataset from the simulated source activity. We simulated sources at 100 random vertices on the pial surface, and ran two sets of simulations: one varying the level of noise in the simulated data and the other varying the magnitude of co-registration error.

Typical per-trial SNR levels for MEG data range from −40 to −20 dB (Goldenholz et al., 2009), and therefore Gaussian white noise was added to the simulated data and scaled in order to yield per-trial amplitude SNR levels (averaged over all sensors) of −50, −40, −30, −20, −10, or 0 dB to generate synthetic datasets across a range of realistic SNRs. Source reconstruction was performed using 10 different models. The reference model used the original link vectors as dipole orientation priors, and the remaining 9 models used vectors with angular differences from the original link vectors ranging from 7 to 63 degrees (in increments of 7 degrees). The orientation of the 9 additional vectors was determined by taking random points on the edge of a cone defined by the reference vector and the angular distance. In these simulations, the co-registration error was 0mm. Within each SNR level, the free energy metric was compared between each model and the reference model.

Within-session head movement and between-session co-registration error commonly combine to introduce a typical magnitude of ^~^5mm (or more) of uncertainty concerning the relative location of the brain and the MEG sensors in traditional MEG recordings (Adjamian et al., 2004; Gross et al., 2013; Ross et al., 2011; Singh et al., 1997; Stolk et al., 2013; Whalen et al., 2008). To simulate between-session co-registration error, we therefore introduced a linear transformation of the fiducial coil locations in random directions (0mm translation - 0° rotation, 2mm - 2°, 4mm - 4°, 6mm - 6°, 8mm - 8°, or 10mm - 10°) prior to source inversion. As in the SNR simulations, source reconstruction was performed using a reference model with the original link vectors as orientation priors, and 9 models using vectors rotated in random directions with angular differences from the original vectors from 7 to 63 degrees. In these simulations, the per-trial amplitude SNR was set to 0dB. Within each level of co-registration error, we compared the free energy between each model and the reference model.

### Head-cast construction

From an MRI-extracted image of the scalp, a head-cast that fit between the participant’s scalp and the MEG dewar was constructed (Bonaiuto et al., 2018a; Meyer et al., 2017; Troebinger et al., 2014b). Scalp surfaces were first extracted from the T1-weighted MRI scans acquired in the first MRI protocol using SPM12 (http://www.fil.ion.ucl.ac.uk/spm/). This tessellated surface, along with 3D models of fiducial coils placed on the nasion and the left and right pre-auricular points, was used to create a virtual 3D model, which was then placed inside a virtual version of the scanner dewar in order to minimize the distance to the sensors while ensuring that the participant’s vision was not obstructed. The model (including spacing elements and ficudial coil protrusions) was printed using a Zcorp 3D printer (Zprinter 510). The 3D printed model was then placed inside a replica of the MEG dewar and polyurethane foam was poured in between the surfaces to create the participant-specific head-cast. The protrusions in the 3D model for fiducial coils therefore become indentations in the foam head-cast, into which the fiducial coils can be placed scanning. The locations of anatomical landmarks used for co-registration are thus unchanged over repeated scans, allowing combination of data from multiple sessions (Bonaiuto et al., 2018a; Meyer et al., 2017).

### Behavioral task

Participants completed a visually cued action decision making task in which they responded to visual instruction cue projected on a screen by pressing one of two buttons using the index and middle finger of their right hand (Bonaiuto et al., 2018a). After a baseline period of fixation, a random dot kinematogram (RDK) was displayed for 2s with coherent motion either to the left or to the right. Following a delay period, an instruction cue (an arrow pointing either to the left or the right), prompted participants to press either the left or right button. The level of motion coherence in the RDK and the congruence between the RDK motion direction and instruction cue varied from trial to trial, but for the purposes of the present study, we analyzed the main effect of visual stimulus onset and button press responses. For a full description of the paradigm and task structure, see Bonaiuto et al. (2018a).

Each block contained 180 trials in total. Participants completed three blocks per session, and 1–5 sessions on different days, resulting in 540–2700 trials per participant (M = 1822.5, SD = 813.21). The task was implemented in MATLAB (The MathWorks, Inc., Natick, MA) using the Cogent 2000 toolbox (http://www.vislab.ucl.ac.uk/cogent.php).

### MEG acquisition and preprocessing

MEG data were acquired using a 275-channel Canadian Thin Films (CTF) MEG system with superconducting quantum interference device (SQUID)-based axial gradiometers (VSM MedTech, Vancouver, Canada) in a magnetically shielded room. A projector was used to display visual stimuli on a screen (^~^50 cm from the participant), and a button box was used for participant responses. The data collected were digitized continuously at a sampling rate of 1200 Hz. MEG data preprocessing and analyses were performed using SPM12 (http://www.fil.ion.ucl.ac.uk/spm/) using MATLAB R2014a. The data were filtered (5th order Butterworth bandpass filter: 2–100 Hz, Notch filter: 50 Hz) and downsampled to 250Hz. Eye blink artifacts were removed using multiple source eye correction (Berg and Scherg, 1994). Trials were then epoched from 1s before RDK onset to 1.5s after instruction cue onset for analysis of visual responses, and from 2s before the participant’s response to 2s after for analysis of movement-evoked responses. Blocks within each session were merged, and trials whose variance exceeded 2.5 standard deviations from the mean were excluded from analysis. The epoched data were then averaged over trials using robust averaging, a form of general linear modeling (Wager et al., 2005) used to reduce the influence of outliers on the mean by iteratively computing a weighting factor for each sample according to how far it is from the mean. Preprocessing code is available at http://github.com/jbonaiuto/meg-laminar.

### Source reconstruction

Source inversion was performed using the empirical Bayesian beamformer algorithm (EBB; Belardinelli et al., 2012; López et al., 2014) as implemented in SPM12. The source inversion was applied to a 100ms time window, centered on the event of interest (the peak of the simulated signal, 100ms following the onset of visual stimuli, or the button press response). These data were projected into 274 orthogonal spatial (lead field) modes and 4 temporal modes. Singular value decomposition (SVD) was used to reduce the sensor data to 274 orthogonal spatial (lead field) modes, each with 4 temporal modes (weighting the dominant modes of temporal variation across the window). For uninformative priors, the maximum-likelihood solution to the inverse problem is:

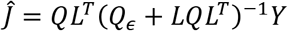

where *ĵ* is the estimated current density across the source space, *Y* is the SVD-reduced measured data, *L* is the lead field matrix that can be computed from the sensor and volume conductor geometry. *Q_ϵ_* is the sensor covariance and Q is the prior estimate of source covariance. We assumed the sensor level covariance (*Q_ϵ_*) to be an identity matrix (see discussion). Most inversion algorithms can be differentiated by the form of *Q* (Friston et al., 2008; López et al., 2014). EBB uses a beamformer prior to estimate the structure of *Q* (Belardinelli et al., 2012; López et al., 2014) based on the sensor-level data:

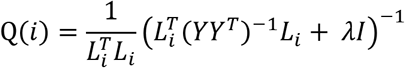

where each element of the diagonal *Q*(*i*) corresponds to a source location /. The lead field of each source location is *L_i_*, ^*T*^ denotes the transpose operator, *I* is an identity matrix, and *λ* is a regularization constant (set to 0). The prior estimates of *Q_ϵ_* and *Q* are then re-scaled or optimally mixed using an expectation maximization scheme (Friston et al., 2008) to give an estimate of *J* that maximizes model evidence. All inversions used a spatial coherence prior (Friston et al., 2008) with a FWHM of 5 mm. We used the Nolte single shell head model (Nolte, 2003).

For MEG source inversion, the accuracy term of the free energy equation is defined as

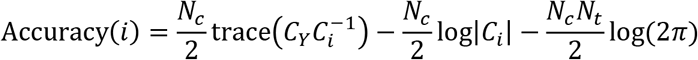

where *N_c_* is the number of channels, *N_t_* is the number of samples, 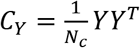 is the data-based sampled covariance, 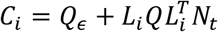 is the model-based sample covariance, and |·| is the matrix determinant operator.

For the EBB algorithm, the complexity term of the free energy equation is dependent on hyperparameters, λ, that control the trade-off between sensor noise *Q_ϵ_* = λ_1_*I_Nc_*, and the beamforming prior *Q* = *λ*_2_*Γ*, where *Γ* is the beamforming prior:

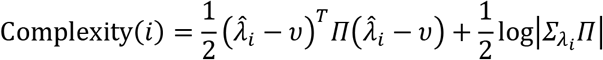

The prior and posterior distributions of λ, *q*(λ*_i_*) and *p*(λ,*_i_*) are assumed to be Gaussian:

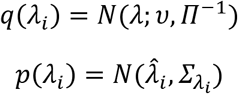

where 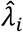 and *Σ_λi_*. are the posterior mean and covariance of the hyperparameters for model *i*. We used non-informative mean and precision (υ and Π) implemented as identity matrices scaled close to zero mean and low precision, as implemented by default in SPM.

## Results

### Different methods for estimating vector orientation yield substantial variation in dipole orientation

We first compared the dipole orientation vectors generated by each of our five methods in terms of the angular difference between vectors at the same vertex on the pial and white matter surfaces, respectively (**Figure 2**). The three methods that utilize only one surface, (downsampled surface normals, cortical patch statistics, original surface normals) generated vectors which were the most similar to each other on both the pial (downsampled surface normals – cortical patch statistics individual subject mean angular difference: 19.16-23.04°; over subjects: M=22.30°, SD=1.51°; downsampled surface normals – original surface normals: 16.00-17.95°; over subjects: M=16.94°, SD=0.67°; cortical patch statistics – original surface normals: 17.80-22.89°; over subjects: M=21.01°, SD=1.55°) and white matter surfaces (downsampled surface normals – cortical patch statistics individual subject mean angular difference: 10.65-11.35°; over subjects: M=10.99°, SD=0.27°; downsampled surface normals – original surface normals: 9.64-10.57°; over subjects: M=10.10°, SD=0.32°; cortical patch statistics – original surface normals: 8.93-9.76°; over subjects: M=9.34°, SD=0.31°). Each single- and multi-surface method generated vectors with mean angular differences from each other of at least 20°, for both the pial surface (downsampled surface normals – link vectors: 22.94-29.56°, M=24.51°, SD=2.15°; downsampled surface normals – variational vector field: 25.52-27.13°, M=26.05°, SD=0.55°; cortical patch statistics – link vectors: 25.26-32.76°, M=27.42°, SD=2.06°; cortical patch statistics – variational vector field: 28.80-32.55°, M=30.91°, SD=1.11°; original surface normals – link vectors: 26.77-33.12°, M=28.44°, SD=2.06°; original surface normals – variational vector field: 26.18-28.43°, M=27.06°, SD=0.75°) and the white matter surface (downsampled surface normals – link vectors: 21.81-29.03°, M=23.46°, SD=2.33°; downsampled surface normals – variational vector field: 27.08-36.26°, M=28.78°, SD=3.06°; cortical patch statistics – link vectors: 22.64-24.82°, M=24.48°, SD=2.34°; cortical patch statistics – variational vector field: 27.31-36.34°, M=29.04°, SD=3.00°; original surface normals – link vectors: 23.38-30.09°, M=25.00°, SD=2.15°; original surface normals – variational vector field: 26.43-35.69°, M=28.31°, SD=3.08°). The two multi-surface methods, link vectors and variational vector field, generated vectors with some of the largest mean angular differences of all method pairs on each surface (pial surface: 67.41-30.74°, M=27.71°, SD=1.40°; white matter surface: 31.33-40.74°, M=33.31°, SD=3.04°). These results were comparable when using surfaces derived from more commonly used 1mm^3^ T1 scans instead of 800μm^3^ MPMs (**Figure S1**). Therefore, rather than being close approximations to each other, each method generates substantially different dipole orientation vectors, even within the multi-surface class of methods.

**Figure 2.**
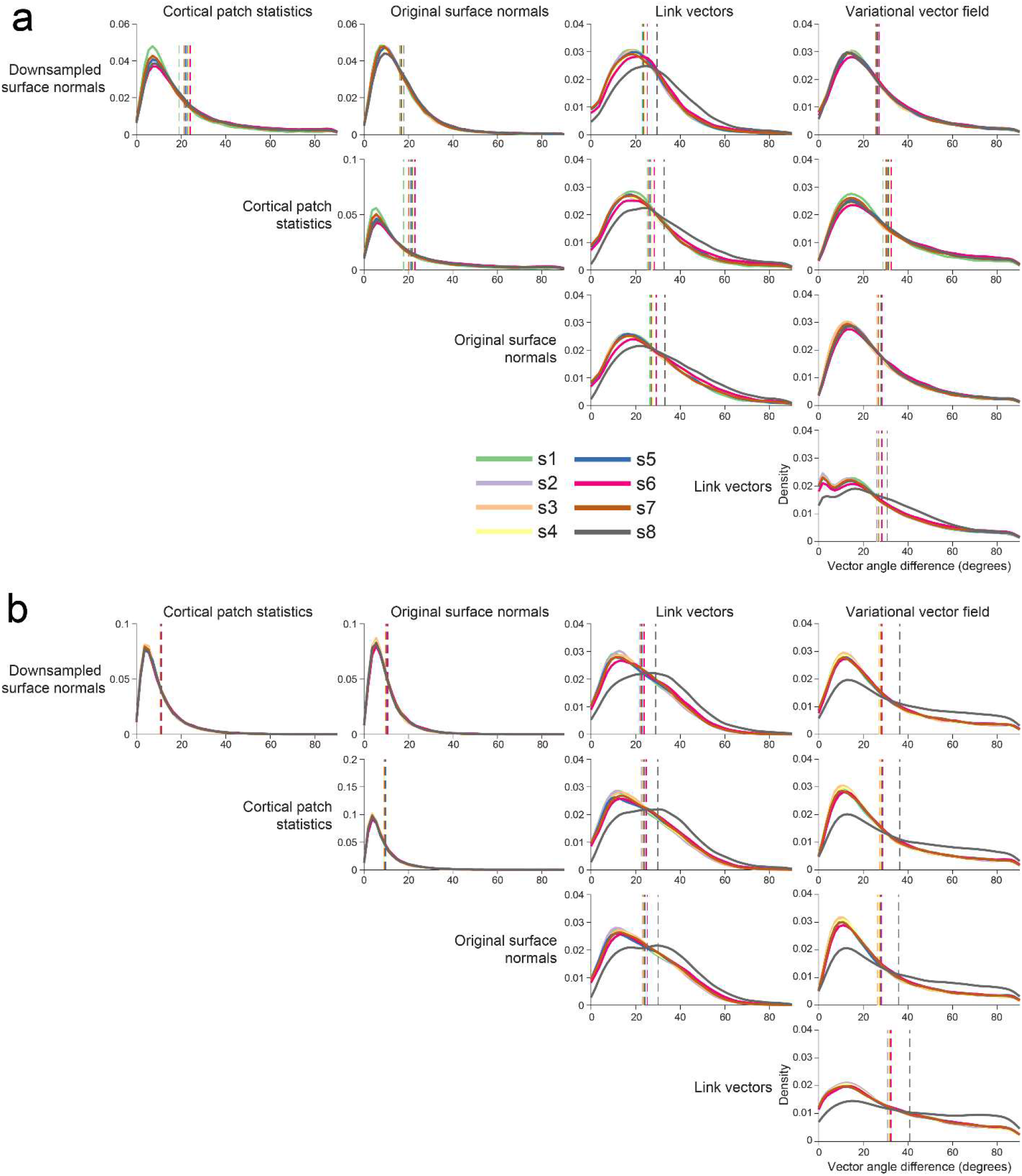
Substantial discrepancy in dipole orientations across methods. Distribution of angular difference between dipole orientations on the pial surface (**a**) and white matter surface (**b**), generated using each method for each participant. Vertical dashed lines show the mean angular difference for each participant.

We next compared dipole orientation vectors generated by each method between the pial and white matter surfaces derived from the 800μm^3^ MPM volumes (**Figure 3**) and 1mm^3^ T1 volumes (**Figure S2**).

Because the link vectors method generates vectors that connect corresponding vertices on the pial and white matter surfaces, the resulting dipole orientations on each surface are equivalent (i.e. the link vector from a particular vertex on the pial surface points in exactly the opposite direction as the link vector from the corresponding white matter surface vertex). All three single-surface methods generated vectors with the lowest average angular difference between pial and white matter surfaces created using either the 800μm^3^ MPM volumes (downsampled surface normals: individual subject mean angular difference=16.88-20.52°, over subjects M=18.23°, SD=1.13°; cortical patch statistics: 22.66-27.81°, M=25.70°, SD=1.67°; original surface normals: 22.90-26.81°, M=24.46°, SD=1.26°) or the 1mm^3^ T1 volumes (downsampled surface normals: 17.89-19.49°, M=18.66°, SD=0.56°; cortical patch statistics: 25.10-27.23°, M=26.23°, SD=0.70°; original surface normals: 24.10-28.80°, M=25.45°, SD=1.46°). The variational vector field method generated dipole orientation vectors that differed the most between the pial and white matter surface (MPM: 37.47-43.54°, M=38.80°, SD=1.99°; T1: 37.07-40.28°, M=39.10°, SD=1.00°). Aside from the link vectors method, there is therefore at least as much variation in dipole orientations between the pial and white matter surfaces within a method as there is between methods for one surface.

**Figure 3.**
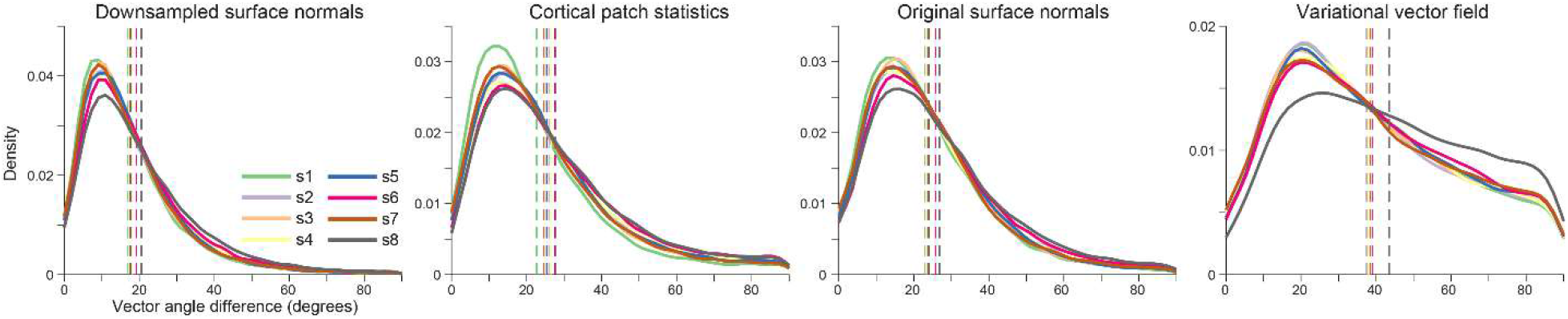
Substantial discrepancy in dipole orientations between pial and white matter surfaces. Distribution of angular difference between dipole orientations at corresponding vertices on the pial and white matter surfaces, generated using the 800 μm^3^ MPM volumes. The link vectors method is not shown because this method generates identical dipole orientations for the pial and white matter surfaces. Each solid line shows the distribution for a single participant. Vertical dashed lines show the mean angular difference for each participant.

### With high precision MEG data, getting the orientation right matters

Having established that each method yields substantially different orientation vectors, we next sought to determine the minimum angular difference between dipole orientations distinguishable by source inversion model comparison, and how this is affected by typical levels of SNR and co-registration error. We therefore simulated dipoles at 100 random source locations on the pial surface and created synthetic datasets with varying SNR and co-registration error levels. We then performed source inversion on the synthetic datasets, using a reference model in which the dipole orientations exactly match those of the simulated dipole, and 9 other models in which the dipole orientations were rotated with respect to simulated dipole orientation. We then compared each of these models to the reference model in terms of the relative free energy, using a significance threshold of ±3 for the free energy difference (indicating that one model is approximately twenty times more likely than the other).

**Figure 4.**
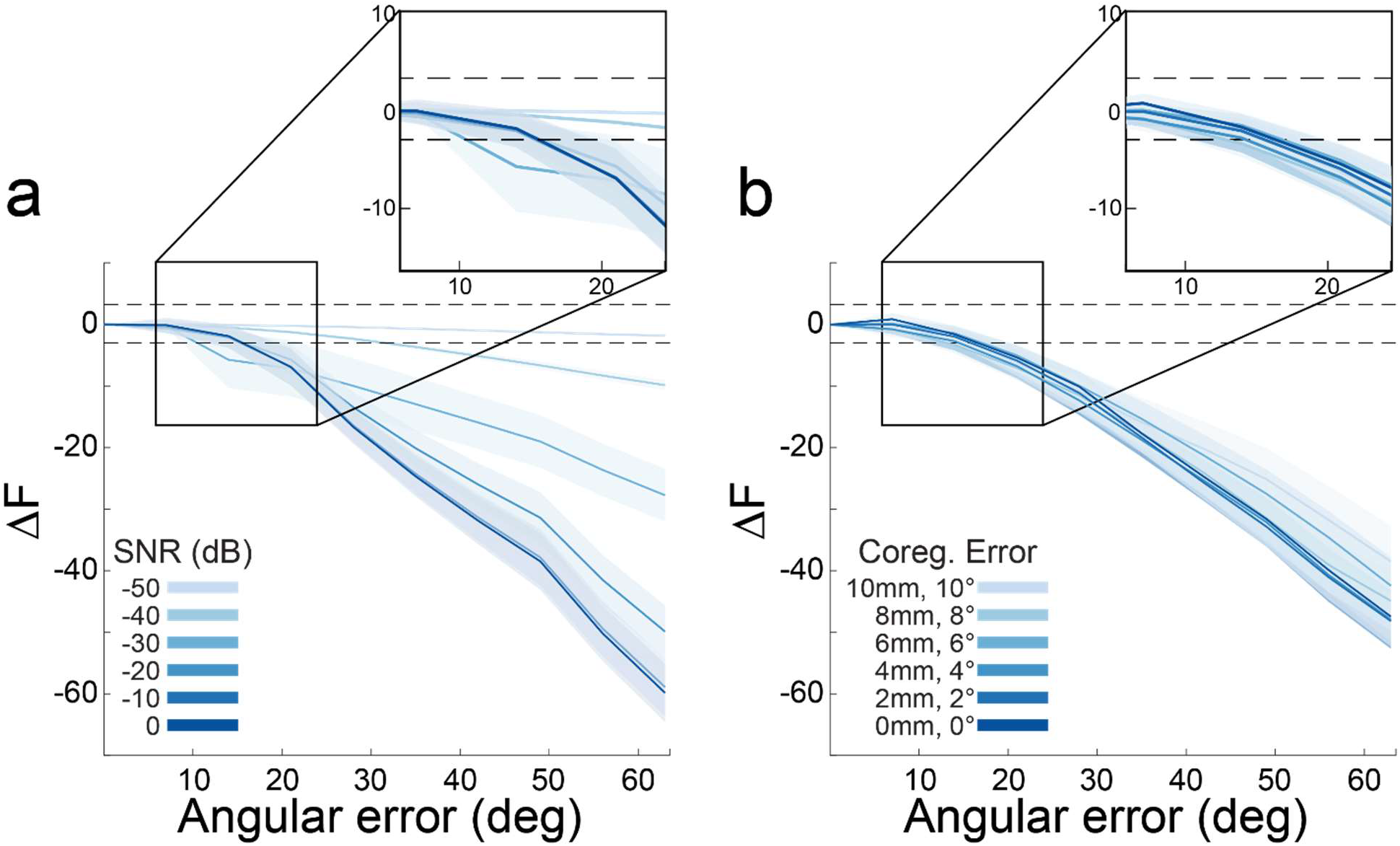
With high precision MEG data, model evidence decreases with dipole orientation error. **a** Each line shows the change in model evidence (ΔF) as the orientation of the dipole used for inversion is rotated away from the true orientation at different SNR levels (co-registration error=0mm). The shaded regions represent the standard error of ΔF over all 100 simulations at each angular error value tested. The lower dotted line (at ΔF=-3) show the point at which the imperfect model is 20 times less likely than the true model. The differences between models become more apparent at higher SNR. **b** As in a, for different magnitudes of co-registration error (SNR=0dB). Co-registration error has a smaller impact than SNR on discriminating between models with different dipole orientations.

At lower SNR levels (−50dB), each model was indistinguishable from the reference model (magnitude of relative free energy less than 3). However, as SNR increased, models with an angular error as low as 15 degrees relative to the reference model started to become differentiable (i.e., a relative free energy difference of >3; **Figure 4a**). Relative model evidence was less dependent on co-registration error, and at all levels tested models with an angular error of at least 15 degrees could be differentiated from the reference model (**Figure 4b**). This angular error is well within the range of the angular differences between vectors generated by each of the methods considered here (**Figure 2**). In other words, given sufficient SNR and co-registration accuracy, one should be able to determine the best method to use with human data based on source inversion model comparison.

### Comparing surface models with empirical head-cast data

We next compared orientation models based on three different evoked responses from human participants. We performed source inversion, and compared the resulting model fits in terms of relative free energy compared to that of the downsampled surface normal model (the current most commonly used method). This was repeated using source space models restricted to the pial surface, white matter surface, and combined pial – white matter surface (Bonaiuto et al., 2018a, 2018b). In this case the combined pial-white model had double the number of sources and these sources could be arranged with identical orientations on each surface (link vectors); or different orientations (cortical patch statistics, downsampled surface normals, original surface normals, and variational vector field).

The evoked response fields (ERFs) were the visually-evoked response to the RDK (visual ERF 1) and instruction cue (visual ERF 2), and the motor-evoked response during the button press (motor ERF). When running the source inversion over the full time course of each ERF, each orientation model yielded slightly different peak cortical locations (**Figure 5**a, b), with the original surface normals and variational vector field methods giving the closest peak coordinates (M=4.88mm, SD=2.93mm), and the cortical patch statistics and link vectors methods yielding coordinates furthest away from each other (M=13.96mm, SD=12.76mm). At each peak location identified by the downsampled surface normals method, the source space ERFs given by the downsampled surface normals, cortical patch statistics, and variational vector field methods, respectively, were most similar to each other, whilst the link vectors methods yielded an ERF with a larger amplitude, and the original surface normals method yielded an ERF with inverted polarity (RMSE<0.1; **Figure 5**c, d). However, at the peak coordinate identified by each method the ERFs were very similar (RMSE<0.05; **Figure 5**e, f).

**Figure 5:**
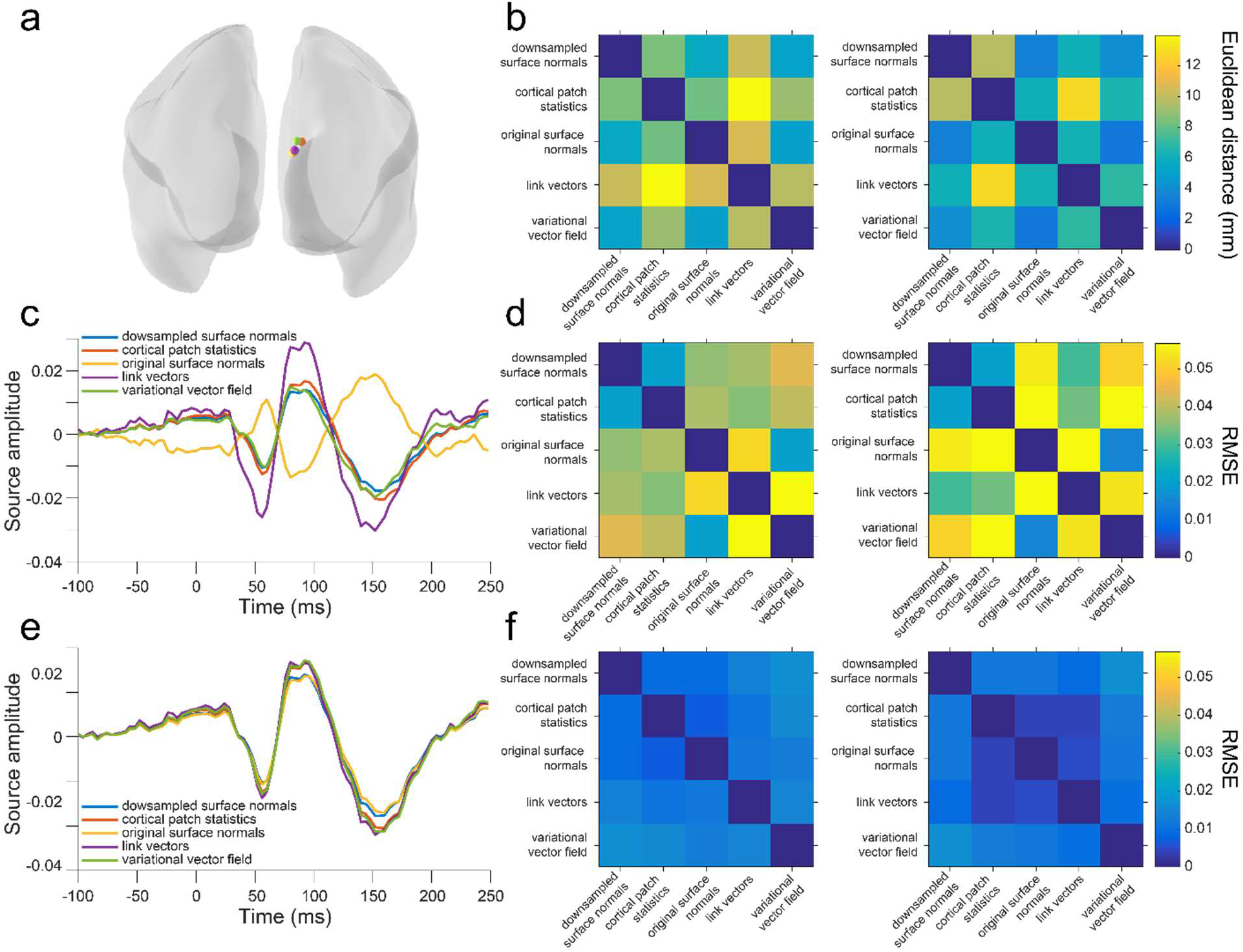
Variation in source localization across methods. **a** Peak source locations on the pial surface for visual ERF1, for each dipole orientation method, in a single subject. **b** Mean (left) and standard deviation (right) of the Euclidean distance between peak source locations for each method, using the pial surface, over subjects and ERFs. **c** Time course of source activity for visual ERF1 for the different methods, at the peak pial source location identified using the downsampled surface normals method, for a single subject. **d** Mean (left) and standard deviation (right) of the RMSE between source activity time courses at the peak source location identified using the downsampled surface normals method, for each method, using the pial surface, over subjects and ERFs. **e** Time course of source activity for visual ERF1 at the peak pial source location identified from each method, for a single subject. **f** Mean (left) and standard deviation (right) of the RMSE between source activity time courses at the peak source location identified using each method with the pial surface, over subjects and ERFs.

We then compared each method in terms of model fit. The link vectors method achieved a significantly better model fit than the downsampled surface normal method in 7/8 subjects for visual ERF 1 and 2 and the motor ERF using the pial surface, 7/8 subjects for visual ERF 1 and the motor ERF and 8/8 subjects for visual ERF 2 using the white matter surface, and 7/8 subjects for visual ERF 1 and 2 and the motor ERF using the two-layer surface (**Figure 6b**). The variational vector field method had significantly better model fit than the downsampled surface normal method in 6/8 subjects for visual ERF 1 and 4/8 subjects for visual ERF2, but only 2/8 subjects for the motor ERF using the pial surface, 1/8 subjects for visual ERF 1 and the motor ERF and 4/8 subjects for visual ERF 2 using the white matter surface, and 5/8 subjects for visual ERF 1 and 2 and 1/8 subjects for the motor ERF using the two-layer surface. The original surface normal method was most similar to the downsampled surface normal method, only being significantly better in 4/8 subjects for visual ERF1, 5/8 subjects for visual ERF2, and 0/8 subjects for the motor ERF using the pial surface, 0/8 subjects for visual ERF 1 and 2 and the motor ERF using the white matter surface, and 1/8 subjects for visual ERF 1, 3/8 subjects for visual ERF2, and 0/8 subjects for the motor ERF using the two-layer surface. The cortical patch statistics method was significantly better than the downsampled surface normals method in 3/8 subjects for visual ERF 1, 6/8 subjects for visual ERF 2, and 2/8 subjects for the motor ERF using the pial surface, 0/8 subjects for visual ERF 1 and 2 and the motor ERF using the white matter surface, and 2/8 subjects for visual ERF 1, 4/8 subjects for visual ERF 2, and 0/8 subjects for the motor ERF.

While the cortical patch statistics and original surface normal methods are an improvement on the widely used downsampled surface normal method, multi-surface methods such as link vectors and variational vector fields achieve better model fits overall, using either single- or two-layer cortical surface models (**Figure 6b**). These results were comparable when using surfaces derived from more commonly used 1mm^3^ T1 scans instead of 800μm^3^ MPMs, with the exception of the variational vector field method, which performed significantly worse than the downsampled surface normal method in 6/8 subjects for the motor ERF using the pial surface (**Figure S3**).

**Figure 6.**
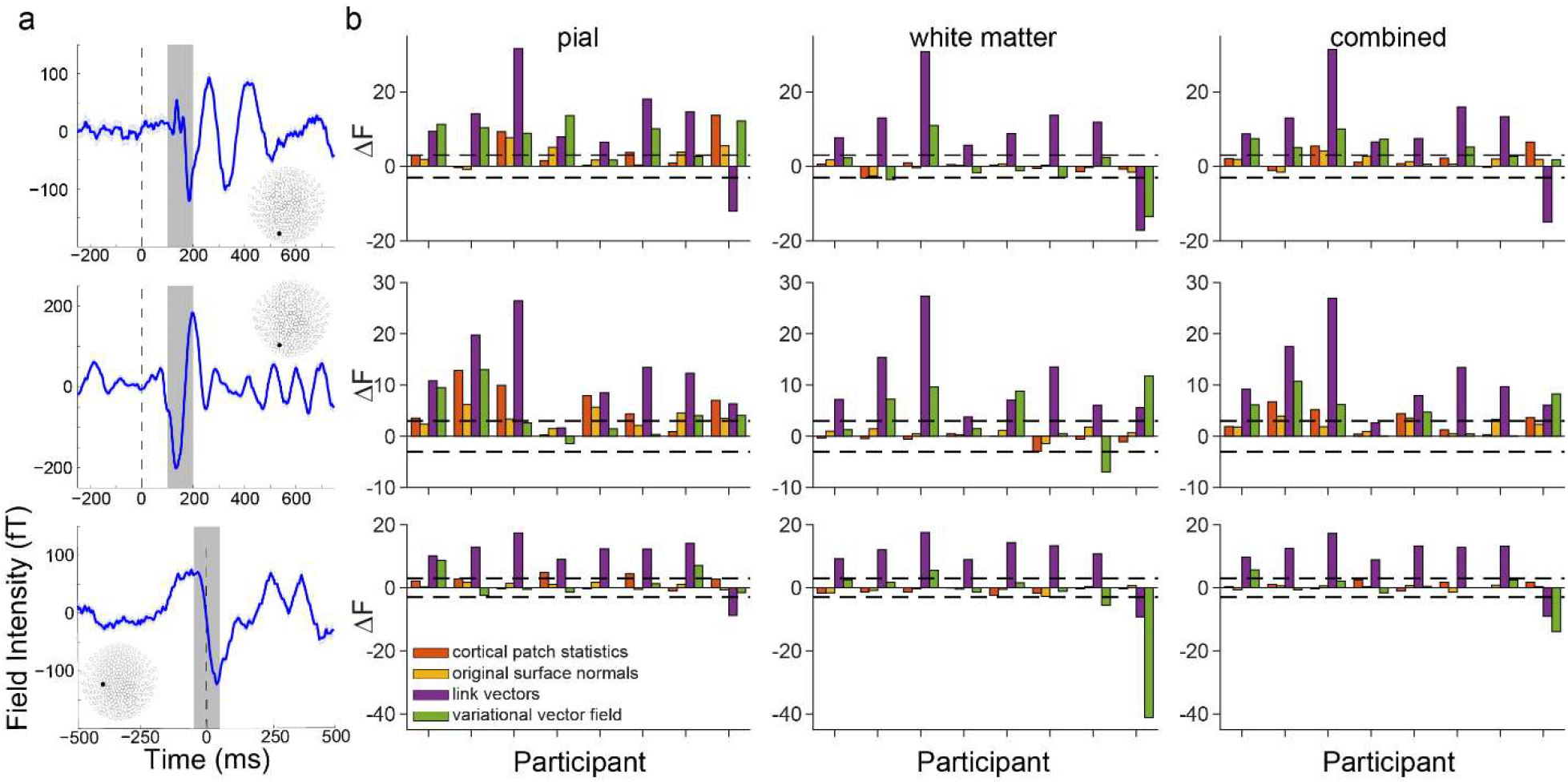
Surface correspondence-based methods yield the best model fit. **a** Trial-averaged event-related fields (ERFs) aligned to the onset of visual stimulus 1 (the random dot kinematogram; top), visual stimulus 2 (the instruction cue; middle), and to the participant’s response (button press; bottom). Data shown are for a single representative participant. The inlays show the MEG sensor layout with filled circles denoting the sensor from which the ERFs are recorded. Each shaded region represents the time window over which source inversion was performed. **b** Change in free energy (relative to the downsampled surface normals model) for each method tested for each participant for visual ERF 1 (top), visual ERF 2 (middle), and the motor ERF (bottom) using vectors derived from 800μm^3^ MPM volumes and source space models based on the pial (left), white matter (center), and combined pial / white matter surfaces (right).

### Interaction between orientation and source space models

We next sought to establish how the orientation models interacted with the different possible choices of source space, specifically the cortical surface used to define source locations and the surface used to compute dipole orientations. We fit the empirical evoked response data, using source space location models based on the white matter, pial, or combined white matter/pial surfaces, and for each orientation model using the white matter, pial, or combined white matter/pial surfaces (with the exception of the combined surface source space orientation model which can only be used with the combined surface source space location model).

**Figure 7.**
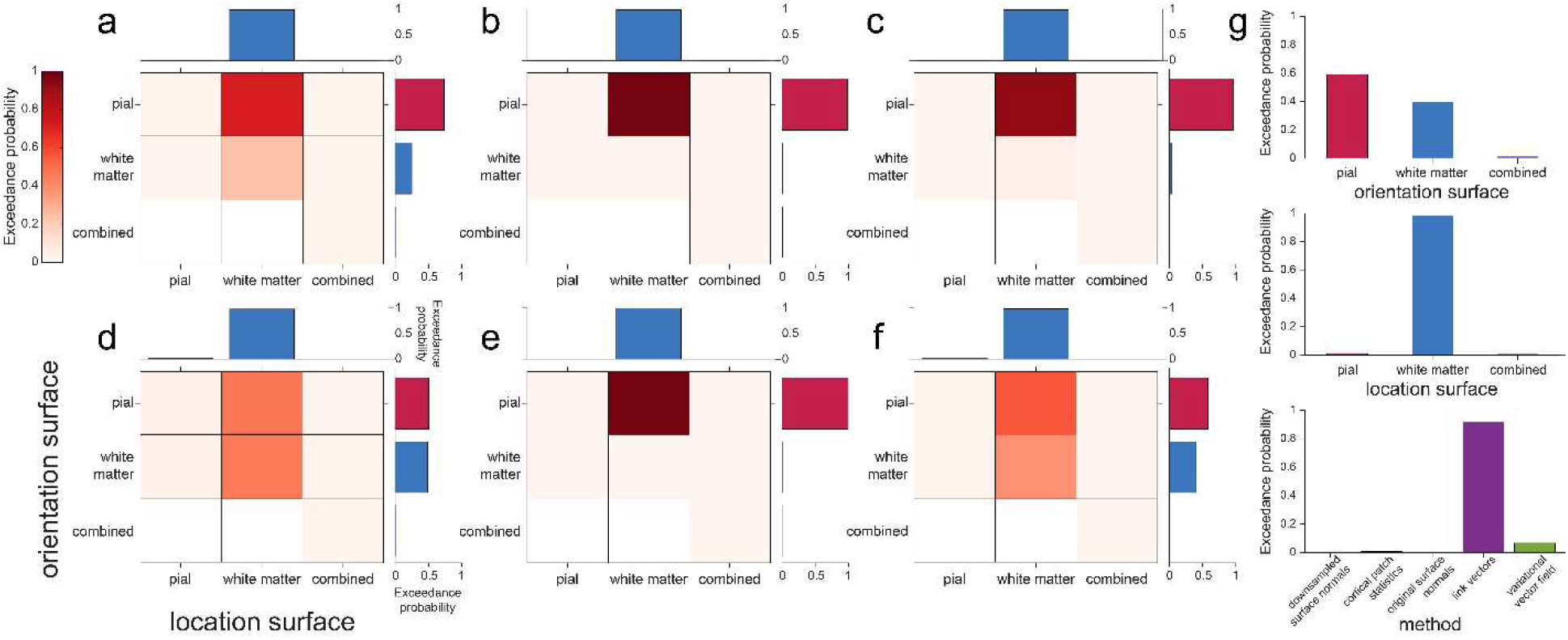
Source inversion using the pial surface and surface correspondence-based methods yield the best model fit overall. **a**-**e** Exceedance probabilities for each combination of source space orientation (pial, white matter, and combined) and location (pial, white matter, and combined) models for each dipole orientation vector method tested (**a** downsampled surface normals, **b** cortical patch statistics, **c** original surface normals, **d** link vectors, **e** variational vector field) using surfaces derived from 800μm^3^ MPM volumes. In each panel the top and right plots show exceedance probabilities for models grouped by source space location or orientation model alone. **f** As in **a**-**e**, for each source space orientation and location models over all dipole orientation vector methods. **g** Exceedance probabilities for each source space orientation model over all source space location models and dipole orientation vector methods (top), for each source space location model over all source space orientation models and dipole orientation vector methods (middle), and for each dipole orientation vector method over all source space orientation and location models (bottom).

To compare source space surface location and orientation models, we used random effects family level Bayesian inference (Penny et al., 2010) over the results from visual ERF 1 and 2 and the motor ERF in all participants. This method groups models into ‘families’, and then combines the evidence of models from the same family to compute the exceedance probability (EP) for each model family. This corresponds to the belief that a particular model family is more likely than the other model families tested, given the data from all participants. We first compared combinations of source space location and orientation surface models within each method, and found that for most dipole orientation methods (downsampled surface normals, cortical patch statistics, original surface normals, and variational vector field), the best source orientation space model was the pial surface and the best source space location model was the white matter surface (downsampled surface normals EP = 0.708, cortical patch statistics EP = 0.968, original surface normals EP = 0.924, variational vector field EP = 0.975; **Figure 7a-c,e**). For the link vectors method, the best source space location model was the white matter surface, and the pial and white matter source space orientation models were nearly indistinguishable (pial orientation model EP = 0.458, white matter orientation model EP = 0.468; **Figure 7d**). This was not unexpected because the orientation vectors generated from these surfaces using the link vectors method are exactly 180° to each other. Over all dipole orientation methods, the best source space orientation surface was the pial surface and the best source space location model was the white matter surface (EP = 0.654; **Figure 7f**). We then grouped models into families based on the source space surface used for orientation and location. This confirmed that the models using the pial surface to compute dipole orientation provided the best model fit, with an EP of 0.608 (white matter surface EP = 0.379), and models using the white matter surface to define source space locations outperformed the others, with an EP of 0.985 (**Figure 7g**). Finally, we grouped models based on the method used to compute dipole orientation vectors and confirmed that the link vectors and variational vector field methods were the best overall, with EPs of 0.918 and 0.064, respectively (**Figure 7g**). Using surfaces obtained from 1mm^3^ T1 volumes yielded the same results (**Figure S4**).

## Discussion

In this paper we show that methods for computing dipole orientation which are based on establishing correspondences between white matter and pial cortical surfaces dramatically outperform methods based on the geometry of a single cortical surface in fitting evoked visual and motor responses. To this end, we compared five different approaches for estimating dipole vector orientation, both in simulations and visual and motor evoked MEG responses.

Our results show substantial variation in dipole vector orientation across the different methods. This indicates that the choice of method is likely to significantly impact the quality of source estimation. At low SNR levels and with head movements commonly observed in conventional MEG recordings, this influence is small or non-detectable. However, with the increased SNR and reduced head movements afforded by high-precision MEG (Meyer et al., 2017; Troebinger et al., 2014b), these differences become distinguishable when average angular errors between methods vary by around 15 degrees. These small orientation errors put a hard limit on any possible improvements in non-invasive estimates of cortical current flow. For example Hillebrand & Barnes (2003) showed that small orientation errors resulted in localization errors which increased monotonically with SNR.

This means that with higher precision MEG recordings, accurate estimation of the dipole orientation becomes increasingly important. Consequently, conventional approaches which estimate vector orientation from a single (downsampled) surface, and based on lower resolution MRI volumes, are likely to offer limited accuracy in source estimation, at least for evoked fields, as analyzed here. By contrast, methods that utilize link vectors between pial and white matter surfaces constructed from higher resolution structural images perform significantly better in explaining observed evoked responses.

The average angular differences between the five methods compared here were substantial, with means of 18-30 degrees, both in high-resolution MPMs and in commonly used T1-weighted structural images with 1mm^3^ spatial resolution. We do not know the ground truth of current flow orientation in the brain, but we show here that the average angular difference between methods is within the range distinguishable in simulated data with SNR and co-registration error levels achievable with high precision MEG. We were therefore able to compare these methods in terms of how well they fit human MEG data, leveraging the free energy metric, in order to determine which method best estimates true dipole orientations.

We were surprised by the large variation in orientation estimates from the same anatomy using different methods. The typical expected orientation differences between methods was ^~^20-30 degrees. This in turn led to differences in estimated source location of ^~^5-14mm. In this study we sought to minimize head-movement and co-registration errors by using head-casts, but in typical MEG studies such additional errors will only add to this variation. Based on these estimates it would seem that if precise anatomical information (e.g. from high resolution MRI volumes) is not available then an approach using some form of loose orientation constraint is advisable (Lin et al., 2006). However, one advantage of being able to exploit anatomical information is to use the sensitivity of MEG to cortical orientation to refine the source localization.

While the family of source space location models based on the white matter surface yielded the highest exceedance probability, the results of the surface comparison varied by evoked response and dipole orientation computation method. Evoked responses can be broken down into temporally dynamic components and therefore may be the result of a complex temporal pattern of signals in both deep and superficial cortical layers. We here used the same 100ms time window for source inversion in all participants and therefore this analysis did not take into account between-participant differences in the timing of evoked responses and could not track the time course of laminar activity. The inherent differences between induced and evoked responses may therefore explain the more variable attribution of the evoked response to pial and white matters surfaces, compared to the bias of high- and low frequency signals towards deep and superficial cortical laminae, respectively (Bonaiuto et al., 2018a). Future extensions of this work could utilize source inversion in successive time bins to address this limitation and generate temporally resolved estimates of laminar activity.

We assumed the sensor level covariance matrix to be diagonal. However, an independent sensor dataset recorded during a similar time period in an empty room, showed off-diagonal structure (**Figure S5**). Importantly, the same pattern of model comparison results was obtained when using a sensor covariance matrix based on these noise measurements (**Figure S6, S7**).

In this work, we used free energy as our metric of model fit but we would expect these findings to generalize across other metrics. For example, we have previously shown that for model comparison problems of the type utilized in this study, free energy is very highly correlated with nonparametric cross validation error measures of model fit (Bonaiuto et al., 2018b).

The present findings do not just impact high SNR MEG recordings obtained with cryogenic sensors, but also for new generations of cryogen-free MEG sensors (optically-pumped magnetometers; OPMs). These sensors can be worn on the head and permit long-duration recordings without head-to-sensor movement, with accurate knowledge of each sensor’s position with respect to the brain (Boto et al., 2018, 2017; Holmes et al., 2018; Iivanainen et al., 2019, 2017; Knappe et al., 2014). Our results show that source estimation for this type of recordings is likely to benefit from methods that estimate vector orientation based on white matter – pial surface vertex correspondences, as opposed to more commonly used techniques employing a single surface.

We here assumed that straight vectors provide the best approximation of the orientation of cortical columns that generate MEG data. However, cortical columns are often curved (Bok, 1929). In future work, the curvature of cortical columns could be approximated using sequences of straight vectors computed from laminar equivolumetric surfaces (Waehnert et al., 2014; Wagstyl et al., 2018). If each vector was tangential to the corresponding segment of the actual (curved) cortical column, this would result in a piecewise linear estimate of column shape, which may allow more precise source localization (Bonaiuto et al., 2018b, 2018a; Troebinger et al., 2014a). This development would benefit from higher resolution (e.g. 7 Tesla) MRI scans, as well as cytoarchitectonic data from histological sections (Amunts et al., 2013; Wagstyl et al., 2018). The current paper provides a novel framework and set of baselines for *in vivo* evaluation of the impact of future columnar models on source modeling.

Our results are likely to impact other methods which require accurate estimation of cortical surfaces and the orientation of surface normal vectors. For example, current flow modelling techniques that estimate the distribution of current delivered with non-invasive brain stimulation approaches such as transcranial direct current stimulation (tDCS; Bestmann and Walsh, 2017; Bestmann and Ward, 2017) estimate the normal component of the electric field across the cortical surface, and relate this component to the observed physiological changes elicited by tDCS (e.g. Laakso et al., 2019; Seo and Jun, 2019). We expect that improved surface segmentation approaches and vector estimation, as introduced in our present study, will provide more accurate estimates of these normal components. This will be relevant for explaining how current delivery via tDCS impacts on physiological and behavioral responses, and whether the normal component of the electric field is indeed important to explain these effects.

## Conclusion

Based on the results of our model comparisons, we have shown that, for evoked responses, source inversion using source locations on the white matter surface and dipole orientation priors computed using link vectors outperforms the other source location and orientation computation methods we tested. We therefore recommend that this approach be used as the default in source inversion.

## Acknowledgements

FA, MF, and MB were supported by the European Union’s Seventh Framework Programme (FP7/2007–2013)/ERC starting grant agreement number 716862 to M. Bonnefond. KW was supported by the Wellcome Trust (215901/Z/19/Z). The Wellcome Centre for Human Neuroimaging is supported by a strategic award from Wellcome (091593/Z/10/Z). The funders had no role in the preparation of the manuscript.

