## Supplemental Material for "Estimates of cortical column orientation improve MEG source inversion"

### Supplementary Material

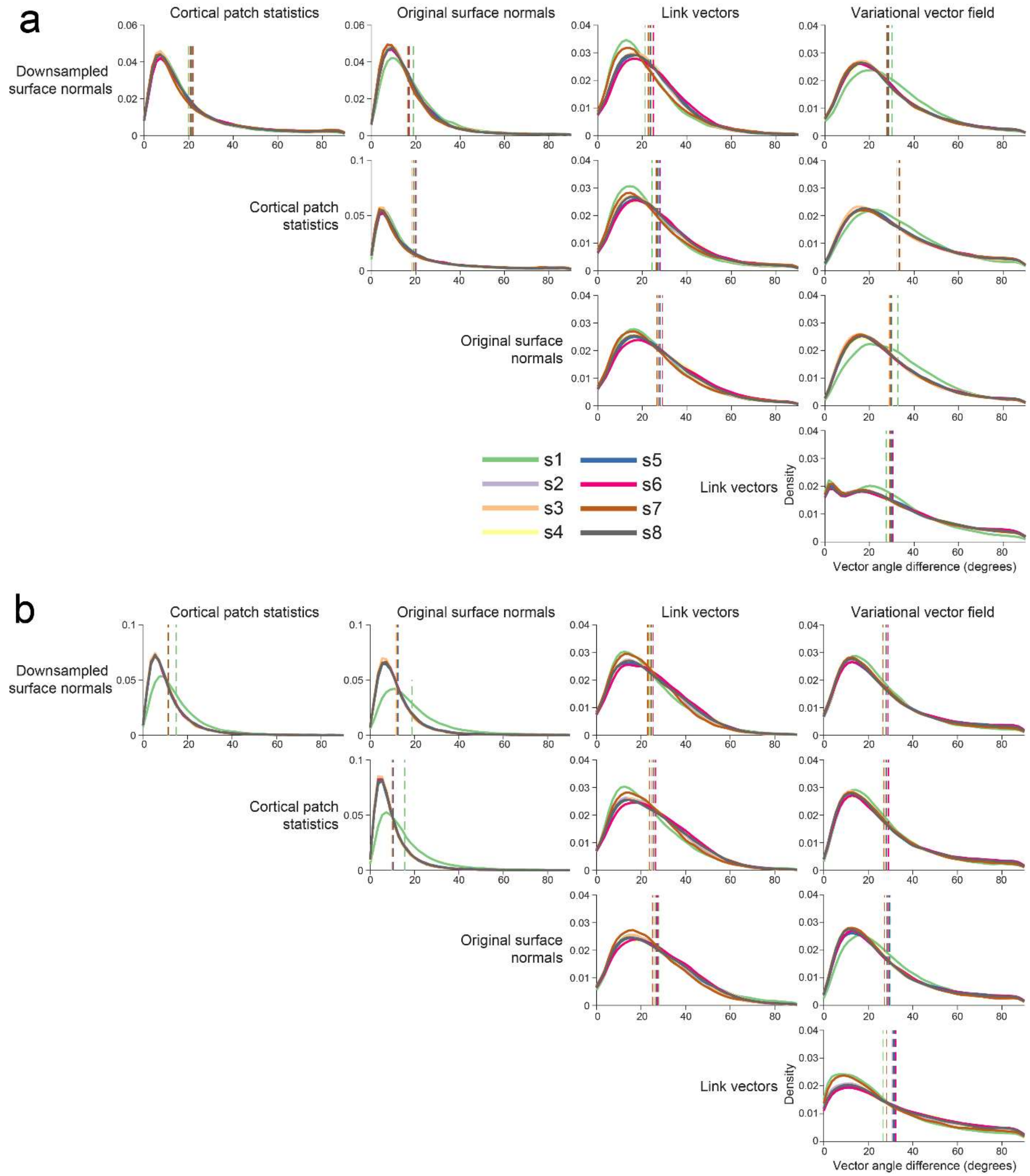


***Figure S1. Dipole orientations across methods using lower resolution scans showed similar patterns of discrepancy as those obtained using high resolution scans.***

***a*** *Distribution of angular difference between dipole orientations generated using each method for each participant using pial surfaces extracted from 1mm$^3$ T1 MRIs. Vertical dashed lines show the mean angular difference for each participant.*

***b*** *As in (a) using white matter surfaces extracted from 1mm$^3$ T1 MRIs.*

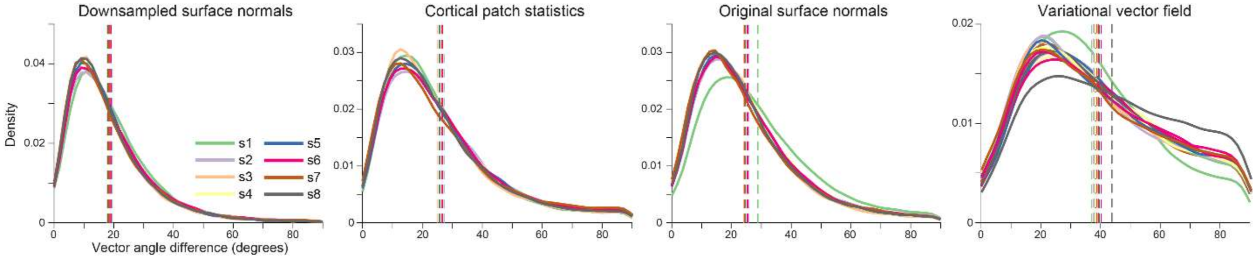


***Figure S2. Substantial discrepancy in dipole orientations between pial and white matter surfaces.*** *Distribution of angular difference between dipole orientations at corresponding vertices on the pial and white matter surfaces, generated using the 1mm$^3$ T1 volumes. The link vectors method is not shown because this method generates identical dipole orientations for the pial and white matter surfaces. Each solid line shows the distribution for a single participant. Vertical dashed lines show the mean angular difference for each participant.*

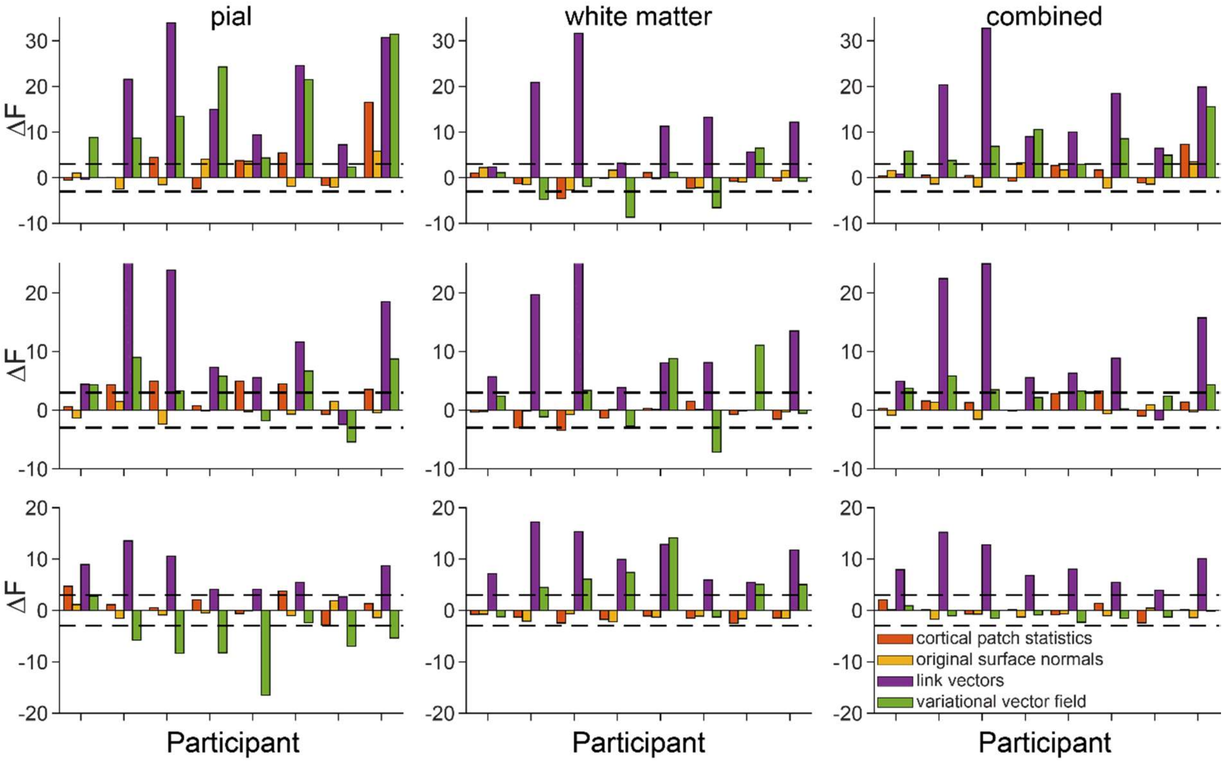


***Figure S3. Results with surfaces from lower resolution scans were comparable to those obtained with high resolution surfaces.*** *Change in free energy (relative to the downsampled surface normals model) for each method tested for each participant for visual ERF 1 (top), visual ERF 2 (middle), and the motor ERF (bottom) using vectors derived from 1mm$^3$ T1 volumes and source space models based on the pial (left), white matter (center), and combined pial / white matter surfaces (right).*

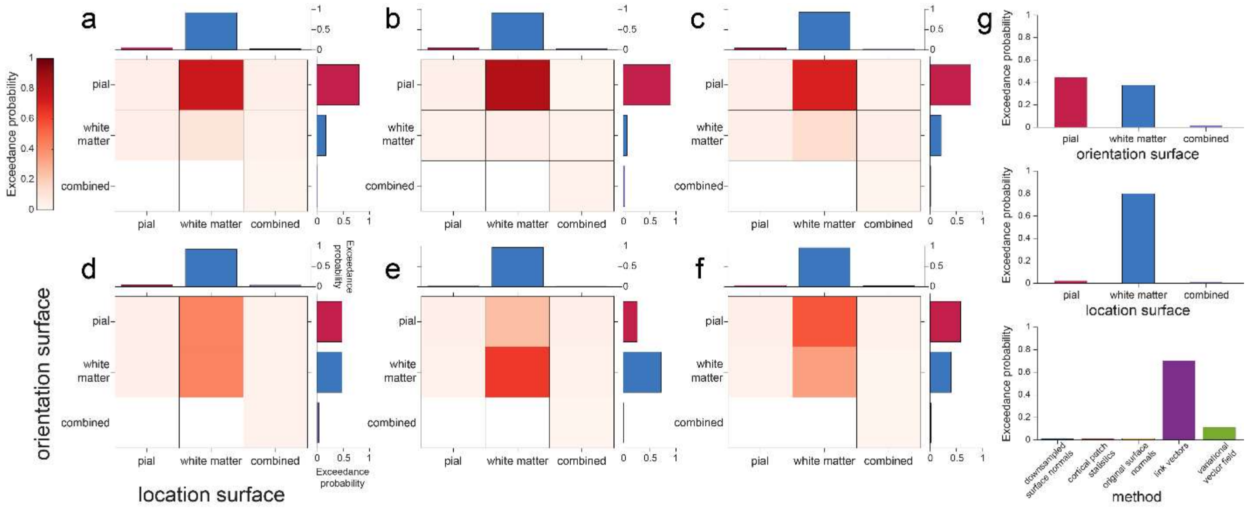


***Figure S4. Results with surfaces from lower resolution scans were comparable to those obtained with high resolution surfaces.*** ***a***-***e*** *Exceedance probabilities for each combination of source space orientation (pial, white matter, and combined) and location (pial, white matter, and combined) models for each dipole orientation vector method tested (**a** downsampled surface normals, **b** cortical patch statistics, **c** original surface normals, **d** link vectors, **e** variational vector field) using surfaces derived from $1mm^3$ T1 volumes. In each panel the top and right plots show exceedance probabilities for models grouped by source space location or orientation model alone.*

***f*** *As in **a**-**e**, for each source space orientation and location models over all dipole orientation vector methods.*

***g*** *Exceedance probabilities for each source space orientation model over all source space location models and dipole orientation vector methods (top), for each source space location model over all source space orientation models and dipole orientation vector methods (middle), and for each dipole orientation vector method over all source space orientation and location models (bottom).*

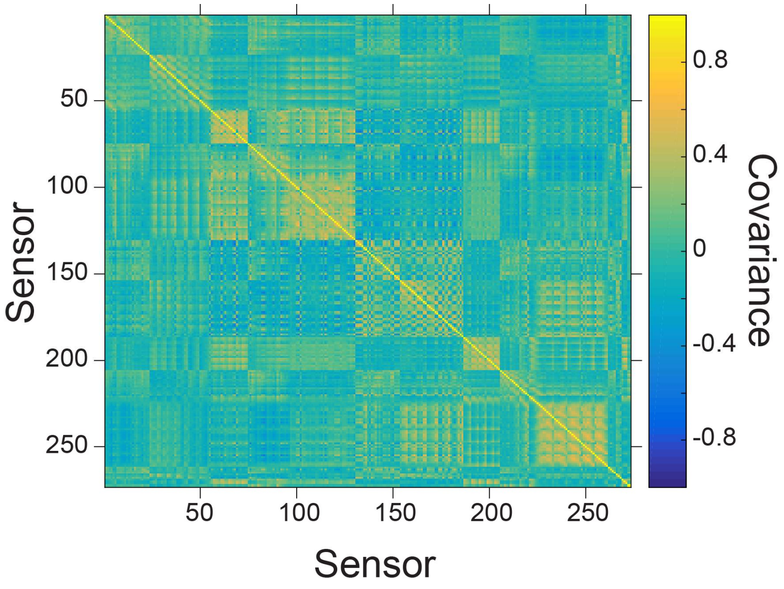


***Figure S5. Sensor activity is correlated.*** *Sensor covariance computed from empty room recordings. The covariance matrix is not perfectly diagonal.*

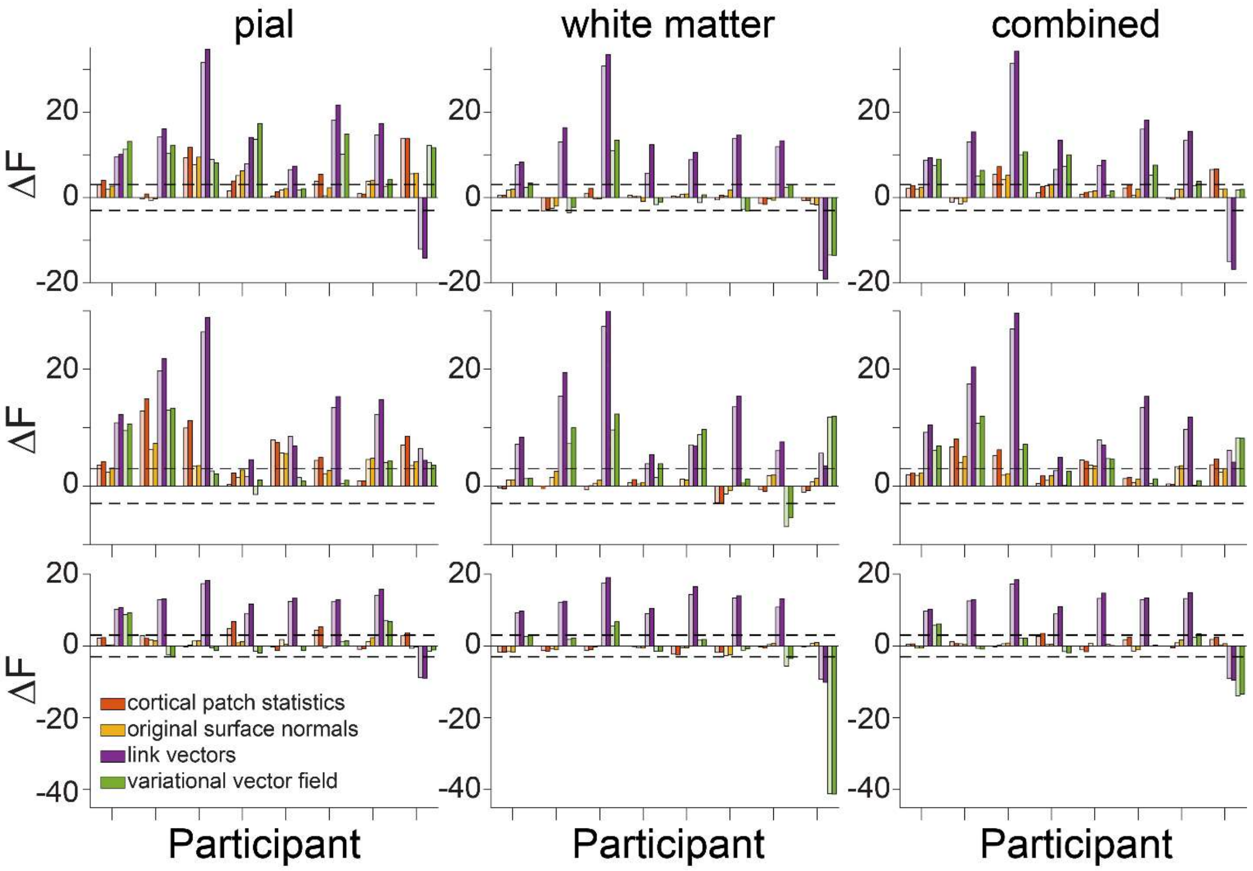


***Figure S6. Model comparison results are not affected by sensor covariance.*** *Change in free energy (relative to the downsampled surface normals model) for each method tested for each participant for visual ERF 1 (top), visual ERF 2 (middle), and the motor ERF (bottom) using vectors derived from 800µm³ MPM volumes and source space models based on the pial (left), white matter (center), and combined pial / white matter surfaces (right). Results obtained assuming an identity sensor covariance matrix are shown in faded colors, and those using a sensor covariance matrix estimated from empty room measurements are shown in bold.*

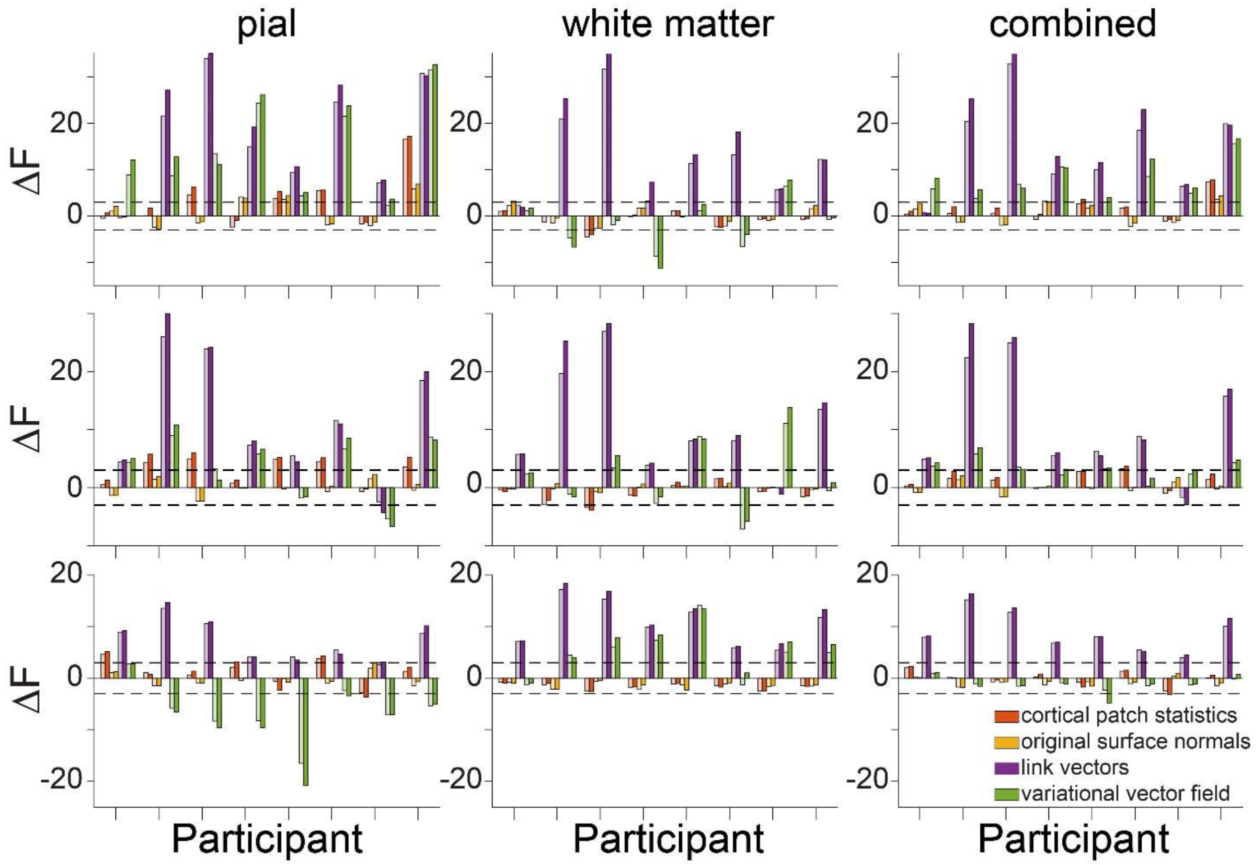


***Figure S7. Model comparison results are not affected by sensor covariance.*** *Change in free energy (relative to the downsampled surface normals model) for each method tested for each participant for visual ERF 1 (top), visual ERF 2 (middle), and the motor ERF (bottom) using vectors derived from 1mm$^3$ T1 volumes and source space models based on the pial (left), white matter (center), and combined pial / white matter surfaces (right). Results obtained assuming an identity sensor covariance matrix are shown in faded colors, and those using a sensor covariance matrix estimated from empty room measurements are shown in bold.*
